# Dispersal in Kentish plovers (*Charadrius alexandrinus*): Adult females perform furthest movements

**DOI:** 10.1101/2023.04.27.538510

**Authors:** Dominic V. Cimiotti, Luke Eberhart-Hertel, Aurélien Audevard, Pere Joan Garcias Salas, Guillaume Gelinaud, Klaus Günther, Afonso Rocha, Rainer Schulz, Jan van der Winden, Heiko Schmaljohann, Clemens Küpper

## Abstract

Dispersal is an important behavioural process that plays a significant role in, among others, speciation, population viability, and individual fitness. Despite progress in avian dispersal research, there are still many knowledge gaps. For example, it is of interest to study how movement propensity (i.e., nomadic vs. philopatric) relates to age- and/or sex-specific patterns of dispersal. Here, we investigated the role of sex and life-stage on natal (i.e., displacement between birth site and first breeding site) and breeding dispersal (i.e., displacement between sequential breeding sites) in the Kentish plover (*Charadrius alexandrinus*). This small and inconspicuous wader is characterised by flexible mating behaviour that includes monogamy, and serial polygyandry. Using a continent-wide dataset of ringing and re-encounter data throughout the species’ range in Europe, we found that adult females generally dispersed further than adult males between seasons, but we detected no sex-difference in natal dispersal distances and no general difference between natal and breeding dispersal distances. Furthermore, females were the main group exhibiting ‘long-distance’ breeding dispersal, which we defined as breeding movements greater than ≥108 km (i.e., upper 10% percentile of our dataset). Our work detected two females breeding in the Mediterranean before dispersing and breeding at the North Sea in the subsequent year, distances of 1,290 and 1,704 km, respectively – this represents the longest known breeding dispersal within the genus *Charadrius*. The long-distance dispersal records we identified are consistent with low genetic differentiation between mainland populations shown in previous work. The dispersive nature of the Kentish plover is likely attributed to its breeding behaviour: polyandrous females exhibit extensive mate searching and habitat prospecting. We recommend that the dispersal traits of Kentish plover be incorporated into the species’ conservation and management planning to more accurately inform models of population connectivity and metapopulation dynamics.

## Introduction

Dispersal plays an important role in population biology as it has many evolutionary consequences for individual fitness and population viability but also adaptation and speciation (e.g., Greenwood 1980, Greenwood & Harvey 1982, D’Urban Jackson et al. 2017). For example, dispersal can increase access to potential mates and reduce competition between relatives or conspecifics (Greenwood 1980). At the population level, dispersal can lead to inbreeding avoidance, range expansion, the colonization of novel habitat, and gene flow throughout a metapopulation (Clobert et al. 2001). Moreover, it is important to understand dispersal dynamics of threatened species so that conservation measures acknowledge a species’ ability to adapt to changing environmental conditions. Here, we study the dispersal dynamics of the Kentish plover (*Charadrius alexandrinus*) and put our findings in the context of other small-bodied avian species with extensive breeding distributions and flexible mating systems – traits that characterize several demographically vulnerable *Charadrius* spp. worldwide.

Dispersal is often categorized by life stage. Natal dispersal refers to the movement of individuals from their birthplace to their first breeding site, whereas breeding dispersal refers to individual movements between successive reproductive attempts (Greenwood & Harvey 1982). Distinguishing between natal and breeding dispersal explains some of the observed variation in dispersal distances within species although a substantial amount of variation in dispersal is also explained by individual state (e.g., sex and previous reproductive success) and habitat quality and availability (e.g., Clark et al. 1997, Skrade & Dinsmore 2010, Rioux et al. 2011, Pearson & Colwell 2014, Swift et al. 2021).

In birds, natal dispersal distances are often larger than breeding dispersal distances (Greenwood & Harvey 1982). For example, Paradis et al. (1998) found larger natal than breeding dispersal distances in 61 of 69 studied British bird species. After having settled as a breeder, site-fidelity can be beneficial in terms of familiarity with former mates, predator avoidance, food resources, and other habitat characteristics of a site (e.g., Greenwood & Harvey 1982), which may explain why breeding dispersal is often less pronounced than natal dispersal. For example, many young colonial seabirds disperse to another colony, but movements of established adult breeders between colonies are rare (e.g., Greenwood & Harvey 1982). However, inbreeding avoidance of young birds might play a key role, too. Alternatively, in nomadic species natal and breeding dispersal are often equally pronounced in all age and sex classes: probably because all birds change sites when habitat conditions deteriorate (e.g., Greenwood & Harvey 1982, Robinson & Oring 1997).

A substantial part of dispersal variation within species is explained by differences between males and females. In mate-competition systems, site familiarity is of greater benefit for the sex that invests more in parental care (e.g., the female), whereas individuals of the other sex may explore different breeding sites in search of potential mates (e.g., Kempenaers & Valcu 2017). In birds, most species show female-biased dispersal as males typically defend territories or resources to attract females. However, in some avian clades, such as Anatidae, males are the more dispersive sex (Greenwood 1980, Clark et al. 1997, Mabry et al. 2013). Male-biased natal dispersal was also found in spotted sandpipers *Actitis macularia*, in which polyandrous females compete over resources (Oring & Lank 1982, Reed & Oring 1993). The mating system itself may be associated with sex-biased dispersal and breeding site fidelity (Kwon et al. 2022), but this relationship is generally weak across birds (Mabry et al. 2013, Trochet et al. 2016).

Despite the aforementioned progress in avian dispersal research, there are still many knowledge gaps – some of which are touched upon in our study. First, it is unclear whether breeding dispersal propensity is high in small-bodied species inhabiting unpredictable coastal and steppe habitats, as this relationship has mainly been documented in larger and more conspicuous species (e.g., Robinson & Oring 1997, Donald et al. 2021). Particularly, it is of interest whether movement propensity (i.e., nomadic vs. philopatric) relates to age- and/or sex-specific patterns of dispersal (e.g., Greenwood & Harvey 1982). Second, dispersal patterns may be even more complex in species with widely spaced breeding distributions and flexible mating systems including serial polygamy with brood desertion and subsequent remating. This behaviour is seen in several *Charadrius* plover species breeding in temperate and tropical latitudes with extended breeding seasons (Eberhart-Phillips 2019, Stenzel & Page 2019). Plover species exhibiting this breeding behaviour are often characterised by male-male resource competition in combination with male-biased brood-care, which may lead to strong sex differences in dispersal. However, patterns may also be population specific due to local constraints such as density dependence and operational sex ratio (Eberhart-Phillips et al. 2018). Third, estimating dispersal rates from restricted study sites may underestimate true dispersal patterns and mislead inference (Barrowclough 1978, Greenwood & Harvey 1982, Paradis et al. 1998). For example, many avian study sites are spatially smaller than the distance that a bird can move within a day (Paradis et al. 1998). Thus, more analyses on broad geographical scale are necessary to cover the full range of dispersal in a species (see Paradis et al. 1998).

To address the aforementioned research gaps in avian dispersal, the Kentish plover (*Charadrius alexandrinus*) serves as an ideal study system. Kentish plovers exhibit a particularly diverse mating system that includes monogamy and serial polygamy with many females deserting early broods soon after hatching to re-mate for another breeding attempt (Rittinghaus 1961, Lessells 1984, Székely & Lessells 1993, Amat et al. 1999, Kosztolányi et al. 2009, Eberhart-Phillips 2019). Additionally, the large breeding range of Kentish plovers includes temperate and subtropical zones of Eurasia and northern Africa where it inhabits coastal and saline inland habitats (BirdLife International 2022). Within Europe, the breeding distribution is scattered along the Atlantic and Mediterranean coasts and inland areas in central and eastern Europe (Keller et al. 2021) with a breeding population of approximately 21,500 – 34,800 pairs (BirdLife International 2015). The species is listed as threatened or declining in many European countries because of degradation and loss of wetland habitat and human disturbance (e.g., destruction of nests) on beaches (e.g., Gomez-Serrano 2020, Thorup & Bregnballe 2021, BirdLife International 2022). Population genetic analyses show that Kentish plovers across continental Eurasia are near panmictic (Küpper et al. 2012, Sadanandan et al. 2019): High gene flow is female-biased and likely driven by serially polyandrous females that are capable of dispersing long distances between breeding attempts (Küpper et al. 2012). However, the genetic data evaluated to-date do not allow for the distinction between natal and breeding dispersal, hindering genetic-based investigations of sex-specific dispersal at different life history stages. Re-encounter data of previously marked birds enable more detailed investigations of dispersal, but previous studies in this species were restricted to local or regional scales (e.g., Székely and Lessels 1993, Foppen et al. 2006).

To characterise the dispersal movements of Kentish plovers in a more comprehensive manner, we utilized a European-wide bird ringing database provided by EURING. We predicted that natal and breeding dispersal distances would not differ in Kentish plovers: dispersal should remain high across both life-history stages due to frequent re-mating in the species and the widely spaced breeding distribution in Europe leading to regular movements of individuals between breeding sites in search of new partners or higher quality habitat. Although both males and females can be serially polygamous, we predicted that breeding dispersal would be higher in females than in males because males acquire nesting territories and invest more in brood care than females, and serial polyandry is more common than serial polygyny in this species (Lessells 1984, Amat et al. 1999, Kosztolányi et al. 2009).

We aimed to use our results gained from the Kentish plover for a comparison with other *Charadrius* plovers as this genus offers a wide range of different breeding systems that might influence the dispersal behaviour.

## Methods

### Source of data and fundamental data categorization

The primary source for our analyses was the ‘EURING Data Bank’ (obtained 9/10/2019; du Feu et al. 2009), which contained 12,057 records of Kentish plovers: 9,651 re-encounters (i.e., re-captures, re-sightings and dead recoveries) of 2,406 individually banded birds. Of those, we excluded entries of individual plovers that were hand-reared or moved before release (13 individuals) and imprecise entries. Imprecise entries were those without a precision of less than one week for the date (35 individuals), or had imprecise geographical locations with an uncertainty of more than 5 km (544 data points of 264 individuals). Finally, we excluded re-encounters of dead plovers excluding those that were found freshly dead (86 individuals).

To separate dispersal from other movements, we classified the status of each ringing or re-encounter data point. We defined ‘certain breeding’ as either being classified as ‘nesting or breeding’ (status ‘N’) or being caught at a nest (catching method ‘N’). We assigned ‘possible breeding’ for all other data with status ‘K’ (‘in colony’) or birds being ringed or re-encountered during the core breeding period between May and June for European populations (see Rittinghaus 1975). As the base data for natal dispersal, we took plovers being ringed as a chick (EURING code ‘1’ for age), because those chicks certainly hatched close to their ringing sites. In total, 5,428 ringing or re-encounter data of 1,906 individuals applied to one of those definitions (i.e., certainly breeding, possibly breeding or ringed as a chick).

We aligned ringing and re-encounter data to identify data that fulfilled one of our breeding status criteria for both the ringing and re-encounter data of an individual. This resulted in 3,223 data points of 1,070 individuals after removing re-encounters of 1^st^ calendar year birds. Consequently, ‘certain dispersal’ refers to cases of ‘certain breeding’ (including data of birds being ringed as chicks) of an individual at the ringing and re-encounter date, whereas the status ‘possible dispersal’ was given for cases with only ‘possible breeding’ either at the ringing or re-encounter stage. We pooled certain and possible breeding dispersal for our analyses to have a larger sample size, but we provide results based on certain dispersal data only if sample sizes were sufficient (e.g., only for between-season breeding dispersal, see below and Fig. S1 in the online supplement).

### Definitions of dispersal types and further data selection

For our analyses, we excluded all dispersal movements within a 10 km radius around the ringing place, and generally dispersal distances up to 10 km, because re-encounters within 10 km were not consistently reported from ringers to national ringing schemes or from national ringing schemes to EURING, and the spatial precision (e.g., data reported on site-level) of most data was too low for fine-scale analyses using the EURING data. Hence, our analyses ignore short distance dispersal movements below 10 km although we acknowledge that such movements may represent a large proportion of ‘dispersal events’ (see Table 1). Here, we focus on medium to long-distance dispersal, which we defined in this study as comprising distances of >10 – 107 km and ≥108 km, respectively. As there is no general threshold for long-distance dispersal in animals and plants (Nathan et al. 2003), and it has therefore to be defined on a case-by-case basis, we classified movements equal or greater than 108 km as long-distance dispersal for our focal group of birds (between-season breeding dispersal) because this threshold represents the upper 10% percentile of our dataset (see Fig. 1b). We did not use the threshold of 50 km used by Stenzel et al. (1994) for long-distance dispersal in the snowy plover because their threshold was case-specific to the US Pacific coast and was not informed by a statistical distribution. We obtained distances between ringing and re-encounter locations directly from the EURING database or measured in QGIS version 3.16.10-Hannover (QGIS Development Team 2020) using its ‘Shape Tools’ extension.

**Table 1:** Summary of published natal and breeding dispersal distances in eight *Charadrius* plover species. Breeding system and brood care information was adopted from Stenzel & Page (2019) and Eberhart-Phillips (2019), data on dispersal distances from Lloyd (2008) and other resources.

| Species | Breeding system (and brood care duties) | Median or mean dispersal distances (km) |  | Maximum dispersal distances (km) |  | Reference for dispersal data<sup>1</sup> |
| --- | --- | --- | --- | --- | --- | --- |
|  |  | natal | breeding | natal | breeding |  |
| Red-breasted plover (*Charadrius obscurus*) | monogamy (biparental) | n.a. | n.a. | 260 | 400 |  |
| Kentish plover (*C. alexandrinus*) | monogamy or serial polygamy (often females and occasionally males desert broods) | n.a. | mean 0.1 – 11.8 (within-season)<sup>2</sup> | n.a. | 170 (f) (within-season) | Székely and Lessels (1993) |
| Kentish plover (*C. alexandrinus*) | see above | median 13.3 | median 1.8 (between-season) | 264 | interval 40-50 (one female 76) (between-season) | Foppen et al. (2006) |
| Kentish plover (*C. alexandrinus*) | see above | median 18 (m), 21 (f); mean 55 (m), 72 (f)<sup>3</sup> | Within-season: median 18.5 (m, f), mean 20 (m), 34 (f)<sup>3</sup>; between-season: median 18 (m), 28 (f), mean 60 (m), 112 (f)<sup>3</sup> | 201 (m), 449 (f) | Within-season: 49 (m), 146 (m); between-season: 1,058 (m), 1,704 (f) | this study |
| Snowy plover (*C. nivosus*) | serial polygamy (females often and males occasionally desert broods) | n.a. | median 170 - 175 (m), 145 - 180 (f) (within- and between-season)<sup>4</sup> | n.a. | 840 (within-season, m), 1,140 (between-season, f) | Stenzel et al. (1994) |
| Snowy plover (*C. nivosus*) | see above | median 4.2 (m), 6.9 (f), 64% <10 km | n.a. | 360 (m), 790 (f) | n.a. | Stenzel et al. (2007) |
| Snowy plover (*C. nivosus*) | see above | n.a. | Within-season<sup>5</sup>: median 0.9 – 1.0 (m), 1.0 – 2.2 (f), 92% <10 km; between-season<sup>6</sup>: median 0.2 - 2,6 (m), 0.3 – 13.0 (f), 85% <10 km | n.a. | n.a. | Pearson & Colwell (2014) |
| Common ringed plover (*C. hiaticula*) | monogamy (biparental) | median 11 | median 0.13 (between-season) | 80 | interval 60-70 (between-season) | Foppen et al. (2006) |
| Piping plover (*C. melodus*) | monogamy (biparental, some females desert broods) | n.a. | mean 35 (m), 26 (f) (between-season)<sup>7</sup> | 1,500 | 595 (between-season) | Haig & Oring (1988) and references cited therein |
| Piping plover (*C. melodus*) | see above | n.a. | 5.8 – 17.8<sup>8</sup> | n.a. | 217 | Rioux et al. (2011) |
| Piping plover (*C. melodus*) | see above | n.a. | n.a. | n.a. | 1,208 (f) (between-season)<sup>9</sup> | Hillman et al. (2012) |
| Piping plover (*C. melodus*) | see above | median 18.6 - 28<sup>10</sup> | median 0.5 - 4<sup>10</sup> | 75 - 306<sup>10</sup> | 71 - 299<sup>10</sup> | Amirault-Langlais et al. (2014) |
| Piping plover (*C. melodus*) | see above | mean 81, median 53, 33% > 100 | mean 23.7, median 0.95; if >50 meters: mean 28.5, median 3.7 (within-season) | 410 | 422 (816) | Swift et al. (2021) |
| Littleringed plover (*C. dubius*) | monogamy (biparental, male > female) | 33 | 5.5 | 250 | 105 |  |
| Littleringed plover (*C. dubius*) | see above | mean 11.1 (m), 12.5 (f) | mean 0.8 (m), 2.2 (f) (between-season) | 333 (f) | 28 (m), 43 (210) (f) (between-season) | Pakanen et al. (2015) |
| Mountain plover (*C. montanus*) | monogamy or simultaneous polyandry (partners have separate clutches and broods) | median 10.6 (m), 7.9 (f); mean 13.0 (m), 10.2 (f)<sup>11</sup> | Within-season: median 0.3 (m), 0.6 (f); mean 2.8 (m), 3.0 (f); between-season: median 0.3 (m), 0.7 (f); mean 2.7 (m), 4.3 (f)<sup>11</sup> | n.a. | n.a. | Skrade & Dinsmore (2010) |
| Eurasian dotterel (*C. morinellus*) | simoultanus polyandry (uniparental male) | n.a. | n.a. | n.a. | 400 |  |

<sup>1</sup> if others than Lloyd 2008 and references cited therein
<sup>2</sup> 0.1 km for pairs moving within a breeding site and 11.8 km if pairs changed site (larger distances in cases of mate change)
<sup>3</sup> only for birds moving >10 km
<sup>4</sup> depending on direction (northwards or southwards), median distances only for birds moving ≥ 50 km
<sup>5</sup> depending on clutch fate (shorter distances after successful breeding)
<sup>6</sup> depending on mate change and clutch fate (shortest distance after successful breeding with same partner)
<sup>7</sup> if not site-faithful; no significant sex difference in breeding dispersal
<sup>8</sup> depending on hatching success (further influence of hatching success of neighbouring pairs)
<sup>9</sup> distance being calculated using Google Earth based on co-ordinates given in the original publication
<sup>10</sup> depending on region
<sup>11</sup> Sex differences in natal and breeding dispersal not statistically significant; longer breeding dispersal after unsuccessful breeding

**Fig. 1:**
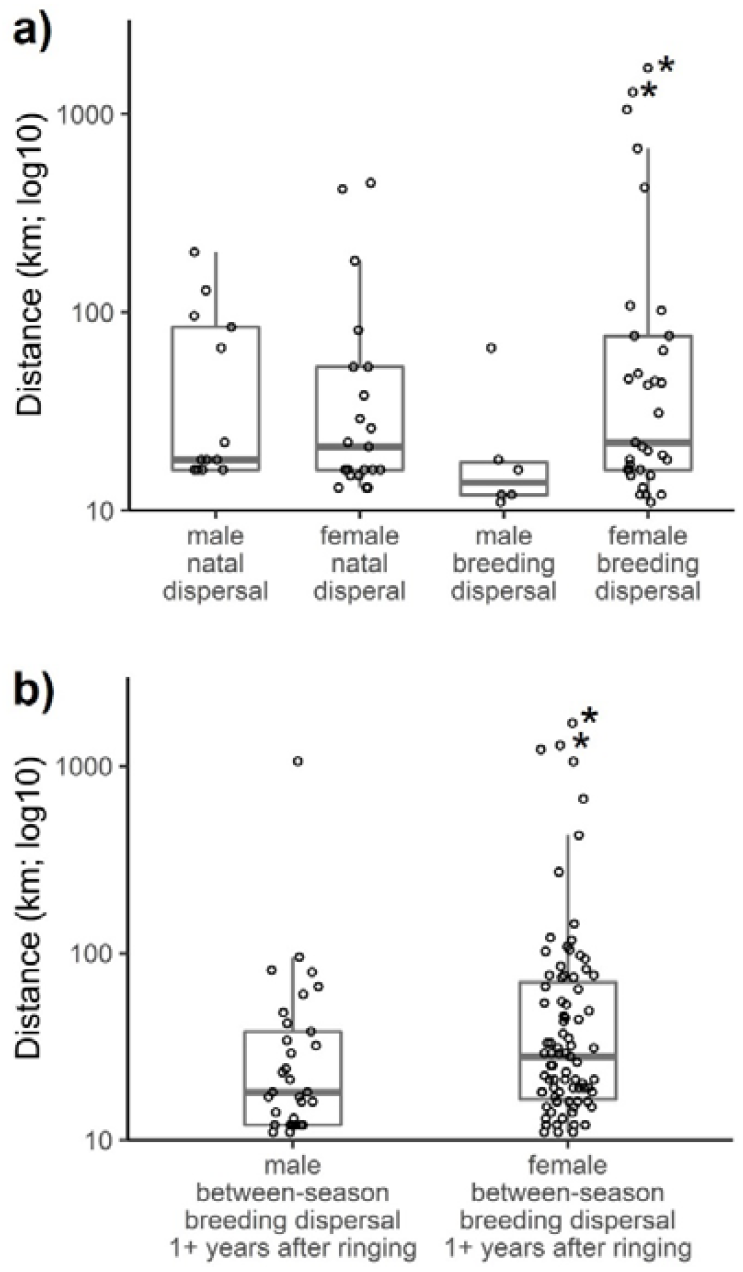
Distribution of sex-specific dispersal distances in European Kentish plovers (*Charadrius alexandrinus*) based on the EURING database: (a) Comparison of all sex-type combinations of natal and breeding dispersal (one data point for each individual within the first year after ringing, see Methods); (b) sex-differences in between-season breeding dispersal without reference to elapsed time since ringing (1 – 13 years). The two recently observed cases of dispersal from the Mediterranean Sea to the North Sea (see text) are indicated by asterisks. Horizontal lines in boxplots show median values, box shows the 25 and 75 percentiles, whiskers the 1.5 times inter-quartile range.

‘Natal dispersal’ referred to as movements by individuals that were ringed as chicks and re-encountered for the first time in their first year after ringing (i.e., the birds’ second calendar year of life, which is the typical age of first breeding in Kentish Plovers, cf. Rittinghaus 1961). Furthermore, the status of their first re-encounter following their hatch year had to be either possible or certain breeding (n = 13 males and 21 females, of which three males and nine females performed certain dispersal). We referred to ‘breeding dispersal’ here for birds that were ringed as adults and re-encountered at least possibly breeding in their first calendar year after ringing (n = six males and 33 females, of which only ten females were categorized as certain dispersal). This made it possible to compare general patterns of ‘natal dispersal’ with ‘breeding dispersal’.

For a more comprehensive analysis of sex differences in between-season breeding dispersal, we included all re-encounter data of an individual without reference to elapsed time since ringing (see Stenzel et al. 1994, Pakanen et al. 2015). Those birds were re-encountered 1 – 13 years after being ringed as an adult (*n* = 33 males, 87 females, of which four males and 22 females referred to certain dispersal). We did so because many individuals were not detected within one year after ringing. However, many of those birds were re-sighted in later years (e.g., over-seen in their old or new breeding site in the meantime). There was no effect of elapsed time since ringing on the observed distances (see Fig. S2 in the online supplement). Only two individuals were detected at more than two places (including the ringing place) >10 km away from each other in different breeding seasons (i.e., those birds performed more than one dispersal movement). For consistency, we used the first observed dispersal movement after ringing for all individuals. Our data included between-season movements of seven birds back to the area around their ringing place (<10 km) following within-season dispersal to another place. We refer to between-season dispersal distances ≥ 108 km as ‘long-distance’ breeding dispersal (i.e., the upper 10% percentile of our dataset).

We refer to within-season breeding dispersal for adult birds that were found at a second place within the same breeding season either in the year of ringing or in subsequent years. However, our sample size was low for this subset of our data (*n* = 20 males, 24 females, of which six females referred to certain dispersal).

### Data analyses

All analyses were performed in R (version 4.1.1, R Core Team 2021). We performed an ANOVA to evaluate sex- and stage-specific differences in (log-transformed) dispersal distances. We performed standard statistical tests (Kruskal-Wallis test, Fisher test) to look for sex differences in between-season breeding dispersal without reference to elapsed time since ringing and in the proportion of long-distance dispersal within the aforementioned categories. We used QGIS (v. 3.16; QGIS Development Team 2020) for visualisation of long-distance breeding dispersal movements.

## Results

### Natal vs. breeding dispersal distances (> 10 km) in the first year after ringing

Median natal dispersal distance of Kentish plovers in the EURING dataset was 18 km (interquartile range (IQR) = 16 – 84 km, maximum = 201 km; *n* = 13) for males and 21 km (IQR = 16 – 53 km, maximum = 449 km; *n* = 21) for females (Fig. 1a). Median breeding dispersal distance was 14 km (IQR = 12 – 17.5 km, maximum = 66 km; *n* = 6) for males and 22 km (IQR = 16 – 76 km, maximum = 1,704 km; *n* = 33) for females (Fig. 1a). We found no effect of sex (*t*1 = -1.70, *p* = 0.09, parameter estimate: - 0.407 ± SE 0.239 for males) and no difference between natal and breeding dispersal (*t*1 = -0.95, *p* = 0.35, parameter estimate: -0.142 ± SE 0.150 for natal dispersal); the sex-type interaction was not significant neither (*t*1 = 1.44, *p* = 0.15, parameter estimate: 0.439 ± SE 0.305 for natal dispersal of males). There were not enough data of certain dispersal to run the same model with those data alone.

### Sex differences in between-season breeding dispersal distances (> 10 km) and the proportion of long-distance breeding dispersal 1+ years after ringing

Males had shorter dispersal distances (median = 18 km, IQR = 12 – 38 km, maximum = 1,058 km; *n* = 33) than females (median = 28 km, IQR = 16.5 – 70 km, maximum = 1,704 km, *n* = 87; Kruskal-Wallis chi-squared = 4.779, *df* = 1, *p* = 0.03; Fig. 1b). This sex difference is consistent if only certain dispersal data are used (see Fig. S1 in the online supplement), but this difference is not statistically significant due to the more limited sample size (*n* = four males, 22 females).The proportion of long-distance breeding dispersal in relation to all observed movements >10 km between season tended to be lower in adult males than in adult females: only one of 33 adult males (3.0%) compared to eleven of 87 adult females (12.6 %) moved ≥108 km, even though the relationship was not significant (Fisher test, *p* = 0.18). Similarly, the proportion of males moving ≥ 108 km in relation to total ringing numbers with given sex (0.2 %, 1 of 403) was about an order of magnitude lower than in females (2.0 %, 11 of 656; Fisher test: *p* = 0.04).

### Extent of long-distance breeding dispersal (≥108 km)

Adult Kentish plovers occasionally moved large distances among breeding sites (Fig. 2). Of the 1,059 adult Kentish plovers that were sexed when ringed, 12 (1.1%) dispersed ≥ 108 km. Dispersal between inland and coastal regions or between different seas (e.g., Atlantic and Mediterranean Sea) was observed in four cases: One male and female certainly dispersed from inland breeding sites in eastern Austria to coastal sites at the Mediterranean Sea in France (1,058 km) or Italy (669 km), respectively, within 1 – 3 years after ringing (Fig. 2). From the Mediterranean Sea, two females certainly dispersed to the North Sea over 1,290 and 1,704 km, respectively, between two successive breeding seasons (Fig. 2).

**Fig. 2:**
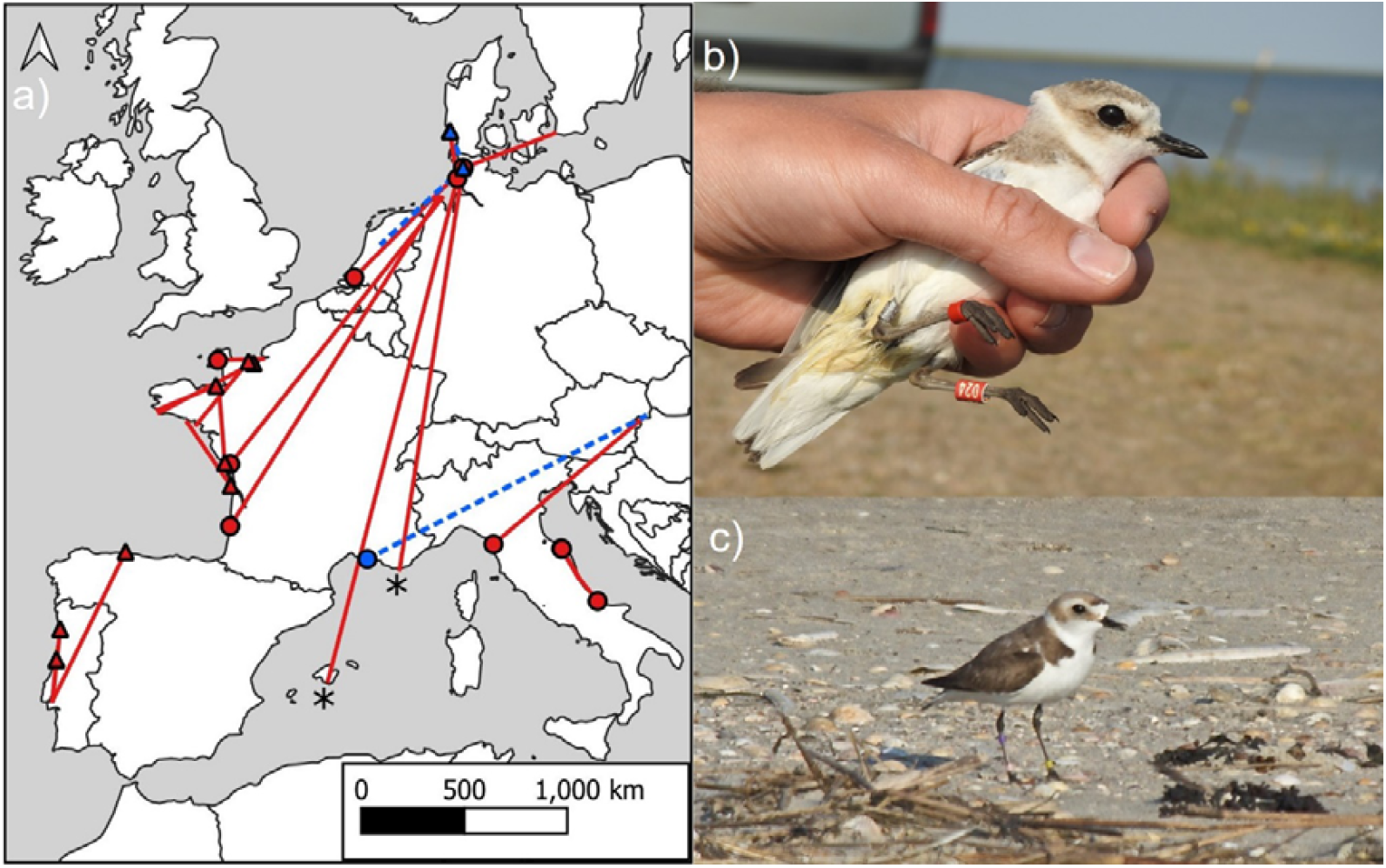
Long-distance breeding dispersal (≥ 108 km) in European Kentish plovers (*Charadrius alexandrinus*): a) Map with data from EURING database (see Methods for details). Places of re-encounters are shown by blue circles (males) and red circles (females). Additional re-encounters from colour-ringing studies in Portugal, France, The Netherlands, and Germany (see Discussion) are supplemented here for visualization with blue triangles (males) and red triangles (females). Independently of data source, the lines connect those places with original ringing places (males: dashed lines, females: solid lines). The ringing places of two females which dispersed from the Mediterranean Sea to the North Sea between the breeding seasons 2017 and 2018 (see panels b and c) are indicated by asterisks. b) Female “Red 024” was ringed during incubation at the salines of Pesquieres near Hyères, France, in 2017 and was re-caught 1,290 km away at its nest in Beltringharder Koog, Germany in 2018 (picture taken by Diana Nett). c) This female dispersed over 1,704 km from the Spanish island of Mallorca to the sandbank of Sankt Peter-Ording, Germany, between the breeding seasons 2017 and 2018 (picture taken by Rainer Schulz).

### Within-season breeding dispersal (>10 km)

Dispersal distances within a breeding season were not significantly different between males (median = 18.5 km, IQR = 17.75 – 19.5 km, maximum = 49 km; *n* = 20) and females (median = 18.5 km, IQR = 12 – 35.75 km, maximum = 146 km, *n* = 24; Kruskal-Wallis chi-squared = 0.009, *df* = 1, *p* = 0.92). The variance of the distances was higher in females than in males, with distances of ≥ 108 km only found in two females. There were not enough data for a separate analysis for the category ‘certain dispersal’ (see Methods), as only data for six females and no data for males were available.

## Discussion

### Natal vs. breeding dispersal in the first year after ringing

Our data show no general difference between natal and breeding dispersal distances (Fig. 1a), indicating that Kentish plovers do not exhibit age-dependent dispersal – a stark contrast to most other bird species, where natal dispersal is more pronounced than breeding dispersal (Greenwood & Harvey 1982, Paradis et al. 1998). Moreover, breeding dispersal at the continental-scale (i.e., > 1,000 km) within the first year after ringing were exclusively detected in adult females (Fig. 1a). However, small scale (< 10 km) dispersal events, which typically comprise the majority of all dispersal movements (cf. Foppen et al. 2006, Skrade & Dinsmore 2014, Pakanen et al. 2015), could not be included in our study due to the limitations of the EURING dataset (see Methods). Consequently, our results are meaningful in regard to medium to long-distance dispersal. This could explain why local-scale dispersal distances of Kentish plovers studied by Foppen et al. (2006) in the Dutch Wadden Sea were further for juveniles than for adults, which contrasts to our results.

A general limitation of large-scale ringing data is that we might have not detected all dispersal movements of an individual, potentially leading to false categorizations (e. g., mixing of early breeding and natal dispersal or within-season and between-season breeding dispersal). Moreover, an observed dispersal movement might be the product of several successive movements. However, as we found no relationship between the time gap and between-season breeding dispersal distance (see Fig. S2), we believe our ringing data reflect true dispersal behaviour of the study species.

### Sex-biased breeding dispersal

Observed dispersal distances were smaller in adult males than in adult females (Fig. 1a, Fig. 1b) and the proportion of dispersing males with long-distance breeding dispersal was nearly ten times lower than the proportion of females compared to ringing numbers (see Results and Fig. 2). To confirm the latter result, we scanned local colour-ringing databanks from Portugal, France, The Netherlands, and Germany (i.e., that were not incorporated into the used EURING dataset). Males only made up two of twelve individuals with between-season long-distance breeding dispersal (≥ 108 km; a male dispersing from The Netherlands to Germany and another male dispersing from Germany to Denmark; see Fig. 2a). In contrast, ten females showed long-distance breeding dispersal, which confirmed the female-bias we found in the EURING data. Therefore, females are the main drivers of breeding dispersal over larger distances in Kentish plovers, which is consistent with the population genetic structure of the species throughout the Palearctic (Küpper et al. 2012).

Female-biased breeding dispersal is likely linked to the mating system of the Kentish Plover, which is dominated by sequential polyandry and reduced parental care in females (e.g., Amat et al. 1999): long-distance dispersal by females could be related to their extensive search for new partners. Because females are rarer than males among breeding Kentish plovers (e. g., 42% in Turkey, Eberhart-Phillips et al. 2018, Eberhart-Phillips 2019), in principle they have the opportunity to re-mate quicker than males (Székely et al. 1999). Long-distance dispersal can enable such females to exploit heterogenous environments and find optimal breeding habitat for the sequential breeding attempts. By contrast, male Kentish plovers need to acquire and defend a breeding territory, with site fidelity providing advantages in local resource competition. Both explanations, female brood desertion for re-mating and male-male resource competition, are not mutually exclusive but likely drive the sex-biased breeding dispersal that we documented.

Despite observing that breeding dispersal was female-biased, we also documented a number of adult males with medium-distance dispersal movements (i.e., >10 – 107 km; Fig. 1a), and one adult male certainly moved >1,000 km within three years after ringing (Fig. 1b; Fig. 2; see Results). Consequently, adult males also make a meaningful contribution to dispersal in this species. Unpaired males, who cannot attract a partner locally, potentially due to male-biased operational sex ratio (i.e., lower mate availability for males; Eberhart-Phillips et al. 2018), may disperse to other sites where the breeding phenology is more favourable to find a suitable mate. Additionally, sometimes Kentish plover pairs have been observed relocating together to an alternative breeding site (Székely and Lessels 1993). Similarly, we observed a breeding pair that moved together over 94 km after nest predation by a crow from a German to a Danish breeding site within one breeding season according to sequential observations of both partners together in three sites (beach of Sank-Peter Ording, Germany; Beltringharder Koog, Germany; island of Rømø, Denmark). Breeding failure has generally been shown to trigger breeding dispersal for both sexes in the Kentish plover (Foppen et al. 2006) and other closely related plover species (e.g., Skrade & Dinsmore 2010, Rioux et al. 2011, Pearson & Colwell 2014).

### High dispersal propensity in the Kentish plover

Despite the above-mentioned sex differences, we detected medium-to-long distance dispersal in all sex-age combinations (Fig. 1a), and some individuals even switched between inland and coastal breeding sites and between the seas (see Fig. 2). Habitat preferences may explain the high dispersal propensity in Kentish plovers. Like the congeneric snowy plover, Kentish plovers breed in highly variable habitats such as saline lakes in steppe environments, on sand banks, in primary dunes, in recently embanked areas, or even in heavily altered and intensely used areas (e.g., salt pans, Rocha et al. 2016) in coastal environments (BirdLife International 2022). Breeding conditions in those habitats can change quickly, for example when temporary salt lakes or lagoons dry out in early summer or storms alter the habitat (Convertino et al. 2011, Cruz-López et al. 2017). Because of the stochastic nature of these breeding habitats, adult Kentish plovers must be flexible throughout their lifetime in order to breed successfully. This may explain the high dispersal potential we documented in our study, which is typical for ecologically similar steppe and coastal birds (e.g., avocets, stilts, and the sociable lapwing *Vanellus gregarius*; Donald et al. 2021, Robinson & Oring 1997, Pigniczki et al. 2019, Joest et al. 2021). Our findings support those of Paradis et al. (1998) that dispersal distances are higher in wetland species and in species with small, scattered, or declining populations.

Additionally, dispersal propensity can be associated with migratory behaviour, as Paradis et al. (1998) found larger dispersal distances in migratory compared to resident bird species. In European Kentish plovers, northern and southern breeding sites are connected by migration (e.g., Bairlein et al. 2014, Spina et al. 2022). Plovers may explore alternative breeding sites during migration or wintering (see Stenzel et al. 1994, Hillman et al. 2021). Conversely, immigrating individuals might affect the migratory composition of populations.

### Genetic consequences

Our findings that Kentish plovers exhibit female-driven long-distance breeding dispersal supports the results of Küpper et al. (2012) showing female-biased gene flow within a near panmictic population throughout continental Eurasia. Küpper et al. (2012) suggested that this female-biased gene flow could be the result of strongly female-biased breeding dispersal and male-biased natal dispersal. This was partially supported by our data, as we observed further breeding dispersal in females than in males. However, we did not detect significant sex differences in natal dispersal. One reason for this difference could be that the data sets are not comparable with respect to sex differences. In particular, the study by Küpper et al. (2012) included many island populations (e.g., from Azores, Madeira, and Cape Verde) that may show different dispersal behaviour due to the closed nature of these isolated populations.

### Comparison with other plover species

Dispersal in Kentish plovers differed from that of other Charadriidae species (Table 1). Mean, median or maximum natal dispersal were greater than breeding dispersal distances in six species of *Charadrius* plovers (see Table 1) and in the northern lapwing (Thompson et al. 1994, Lislevand et al. 2009), another member of the Charadriidae family. The only exception was the red-breasted plover (*C. obscurus*), see Marchant & Higgins 1993 and Table 1. In this context, the observed pattern in our study is remarkable, and the difference to most other studies might be either due to true biological differences between the species or to methodological differences. The latter implies that most studies only examined small population segments, and long-distance dispersal was only anecdotally precepted in most of those studies. In contrast, the EURING dataset covered a large geographical range allowing us to examine the differences between demographic classes and sexes more comprehensively. Thereby, the known maximum breeding dispersal distance of the Kentish plover increased tenfold from 170 km (Székely and Lessels 1993) to 1,704 km.

The observed long-distance dispersal in Kentish plovers includes the farthest recorded breeding dispersal movements within the genus *Charadrius* (see Table 1). Movements between breeding sites of more than 1,000 km were only published for two other species: A female piping plover dispersed from Michigan, USA, over 1,208 km into the range of the Atlantic population in North Carolina, USA (Table 1; Hillman et al. 2021). A female snowy plover from Monterey Bay, California, USA, dispersed 1,140 km northwards to Washington state (Stenzel et al. 1994).

The role of sexual selection on sex-biased dispersal has also been described in the polyandrous snowy plover *C. nivosus* of North America (Table 1; Stenzel et al. 1994, Pearson & Colwell 2014), whereby long-distance breeding dispersal is similarly female-biased. Female-biased breeding dispersal is also present in little ringed plovers *C. dubius* (Pakanen et al. 2015), a species with male-biased brood care and occasional brood desertion by the female (Stenzel & Page 2019). In contrast, no sex differences were found in piping plover *C. melodus* and mountain plover *C. montanus*, two species in which female brood desertion or polyandry is extremely rare (Haig & Oring 1988, Skrade & Dinsmore 2010) – indicating that sex-biased dispersal likely may go hand-in-hand with breeding system variation in *Charadrius* plovers and other shorebirds (Kwon et al. 2022).

### Monitoring implications

The exchange of individuals among breeding sites may occasionally lead to an over-estimation of population sizes, but also morality rates (e.g., quantifying true mortality is complicated by permanent emigration when basing inference on local mark-recapture observations), which should be considered when interpreting local population trends and forecasting population viability (Eberhart-Phillips & Colwell 2014). To avoid double-counts of dispersing individuals, population counts should be synchronised on a geographical scale that acknowledges movement tendencies (Eberhart-Phillips et al. 2016).

## Conclusions

The dispersal propensity of the Kentish plover is high – an encouraging sign that the species can find patchy and far-distant suitable breeding habitats despite human-induced habitat destruction. Hence, conservation efforts such as the maintenance or restoration of suitable habitat should include abandoned or potential breeding areas (e.g., in Great Britain and southern Scandinavia) in contrast to more site-faithful species, for which conservation efforts should be focussed on presently used areas and their surroundings. Even if recently used breeding areas are managed and kept intact for Kentish plovers, alternative breeding sites should remain available for dispersing individuals, especially when conditions worsen elsewhere within the species’ range.

However, it is essential to preserve key breeding areas (e.g., in southern Europe) to maintain source populations (Tittler et al. 2006). Breeding success should be enhanced by conservation measures (e.g., Cimiotti & Hötker 2013) even at productive breeding sites to produce a surplus of young birds for the colonisation of other areas. Potentially misleading interventions such as nest protection measures in areas with low chick survival should be carefully weighted with respect to the high dispersal propensity of the species. The aim should be a network of well-managed breeding sites with a coordinated population monitoring throughout the species’ range. Illustrative cases of exceptional long-distance dispersal, as observed in our study, can be used for raising public awareness for endangered species, and thereby increase public support to respect protection zones for beach-nesting birds.

Our data supports recent findings that high dispersal propensity is intertwined with sex differences in breeding biology (d’Urban-Jackson et al. 2017, Kwon et al. 2022). This is especially true for species that are not classically nomadic: other factors such as the mating system can lead to high dispersal propensity. Therefore, a simple classification into nomadic vs. philopatric species has to be treated with care.

## Acknowledgments

We are grateful to the European Union for Bird Ringing (EURING) which made the recovery data available through the EURING Data Bank and to the many ringers and ringing scheme staff who have gathered and prepared the data. We are grateful to P. Gienapp and D.S. Cimiotti for the statistical advice.

## Funding

The Ministry for Energy Transition, Climate Protection, Environment and Nature of Schleswig-Holstein (Germany) financed a previous study on the species. CK was supported by the Max-Planck Society, LEH was funded by the German Science Foundation (DFG Eigene Stelle grant: EB 590/1-1).

## Data Availability Statement

Data used in the study can be obtained after request at the EURING Data Bank, http://www.euring.org/ (Accessed date 9 October 2019). The additional data used for the Discussion can be obtained from the first author.

## Conflicts of Interest

The authors declare that they have no conflict of interest.

## Notes

### Competing Interest Statement

The authors have declared no competing interest.

