## Supplementary Figures 1 and 2 for "Dispersal in Kentish plovers (*Charadrius alexandrinus*): Adult females perform furthest movements"

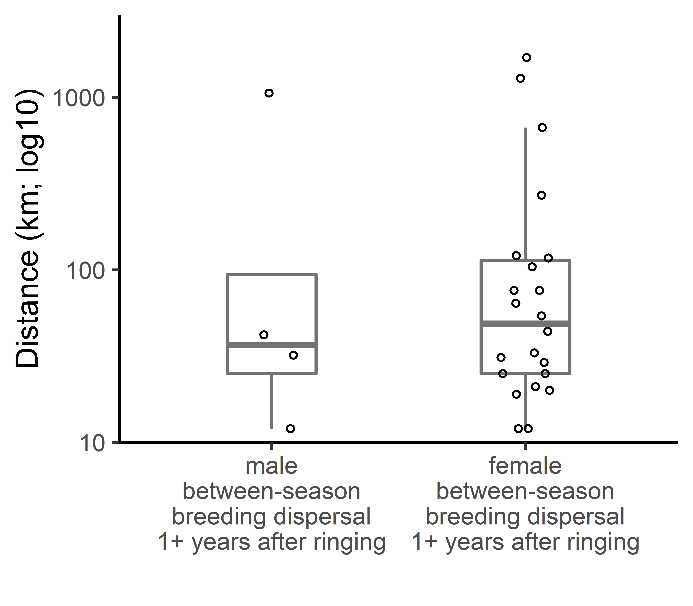

Fig. S1: Sex-differences in between-season breeding dispersal for cases of certain dispersal only.

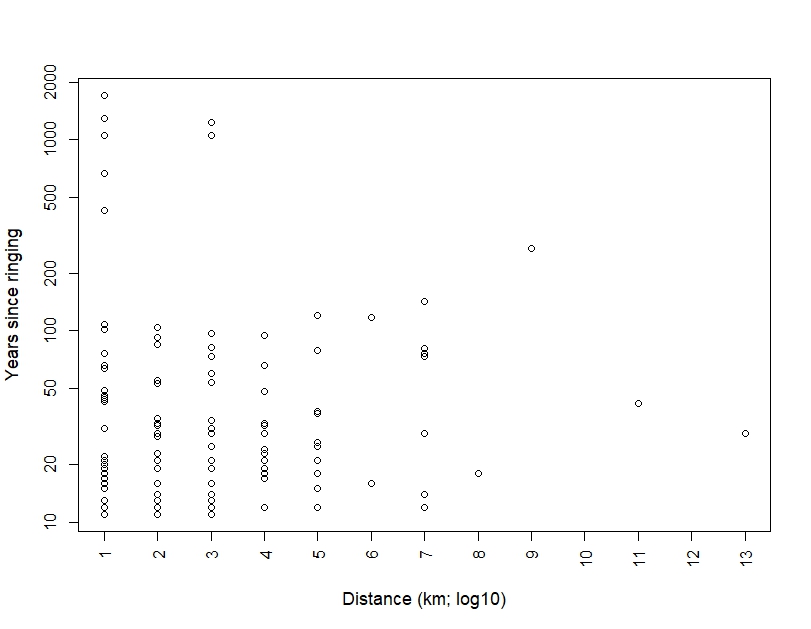

Fig. S2: Between-season dispersal distance in relation to elapsed time since ringing (no effect; LM: *t* = 0.26, *p* = 0.80, parameter estimate: 0.005 ± SE 0.02).
